# The RZZ complex facilitates Mad1 binding to Bub1 ensuring efficient checkpoint signaling

**DOI:** 10.1101/322834

**Authors:** Gang Zhang, Thomas Kruse, Jakob Nilsson

**Affiliations:** The Novo Nordisk Foundation Center for Protein Research, Faculty of Health and Medical Sciences, University of Copenhagen, Blegdamsvej 3B, 2200 Copenhagen, Denmark; Cancer Institute, The Affiliated Hospital of Qingdao University, Qingdao, Shandong 266061, China; Qingdao Cancer Institute, Qingdao, Shandong 266061, China

## Abstract

The recruitment of Mad1 to unattached kinetochores is essential for generating a “wait anaphase” signal during mitosis yet Mad1 localization is poorly understood in mammalian cells. In yeast the Bub1 checkpoint protein is the sole Mad1 receptor but in mammalian cells the Rod-ZW10-Zwilch (RZZ) complex is also required for Mad1 kinetochore localization. The exact function of the two mammalian Mad1 receptors and whether there is any interplay between them is unclear. Here we use CRISPR genome editing to generate RNAi sensitized human cell lines revealing a strong requirement for both Rod and Bub1 in checkpoint signaling. We show that the RZZ complex facilitates Mad1 binding to Bub1 and that a region of Bub1 overlapping the Mad1 binding site stimulates RZZ kinetochore recruitment. The requirement for RZZ in the checkpoint, but not Bub1, can be bypassed by tethering Mad1 to kinetochores or by increasing the strength of the Bub1-Mad1 interaction. Our data support a model in which the primary role of RZZ is to localize Mad1 at kinetochores allowing for the efficient checkpoint generating Mad1-Bub1 interaction. As such, the core checkpoint principle is conserved from yeast to man.

---

In response to unattached kinetochores the spindle assembly checkpoint (SAC) is activated resulting in kinetochore recruitment of Mad1/Mad2, Bub1 and BubR1 checkpoint proteins in a manner stimulated by Mps1 kinase activity^1^. The recruitment of the Mad1/Mad2 complex, mediated by Mad1 kinetochore interactions, is essential because it catalyzes the first step in the generation of the mitotic checkpoint complex (MCC) that is the SAC effector delaying anaphase onset^2–5^. Understanding Mad1 interactions at kinetochores is therefore crucial for understanding the mechanism of SAC activation. In yeast Bub1 is the only receptor for Mad1 and formation of the Mad1-Bub1 complex requires Mps1 phosphorylation of conserved domain 1 (CD1) in Bub1^6–8^. This mode of Mad1-Bub1 interaction is conserved to mammalian cells^4,5,9,10^. However in mammalian cells the RZZ complex also contributes to localization of Mad1 and checkpoint activity (Supplemental Fig. 1)^11–15^. Several conflicting models regarding the specific mechanistic role and relative importance of the two Mad1 receptors in SAC signaling have been proposed ^10,16–20^. This is possibly due to varying degrees of depletion efficiencies and the fact that even minute amounts of checkpoint proteins are able to generate a checkpoint signal^21^. Furthermore whether there is any interplay between the two Mad1 receptors remains unexplored. Here we find a strong requirement for both Rod and Bub1 in generating a checkpoint signal and find that the two Mad1 receptors work in an interrelated fashion to facilitate checkpoint signaling.

To obtain a quantitative comparison of Rod and Bub1 null phenotypes we used CRISPR technology to target exon 2 of Rod and Bub1 in HeLa cells. Extensive screening did not result in any Rod null clones suggesting that Rod is essential in HeLa cells but we did identify clones with reduced expression of Rod, referred to as Rod C (Supplementary Fig. 2a-d). We isolated apparent Bub1 null clones (Bub1 C) as judged by an antibody targeting the N-terminus of the protein while a phospho-specific antibody recognizing S459/T461 phosphorylation (pSpT) detected residual Bub1 at kinetochores (Supplementary Fig. 2e-g). We then used specific RNAi oligoes (Supplemental Fig. 1a-c and ^9^) to further deplete Rod and Bub1 in the respective CRISPR cell lines (referred to as Rod CR and Bub1 CR). This resulted in almost undetectable amounts of protein remaining (Supplemental Fig. 2a,g).

In agreement with the reported role of RZZ and Bub1 in chromosome segregation^21–23^ Rod CR and Bub1 CR exhibited delays in alignment of chromosomes to the metaphase plate and after a delay they often exited with unaligned chromosomes suggesting a weakened SAC (Fig 1a and Table 1). To measure the strength of the SAC, we challenged Rod CR or Bub1 CR with nocodazole or taxol. Here, we observed a strong impairment in checkpoint signaling with only minor SAC activity remaining under both conditions (Fig. 1b,c). Importantly, these effects were fully restored by exogenous RNAi resistant Venus-Rod or Venus Bub1 1-553 attesting to specific removal of the two proteins (Fig. 1b). We note that Mad1 levels in Rod RNAi treated cells and Rod CR are very similar arguing that there is not a direct correlation between Mad1 levels and SAC strength (Supplemental Fig. 2c,d). Moreover, RNAi depletion of Bub1 in Rod CR or Rod in Bub1 CR or their co-depletions in HeLa or U2OS cells resulted in a complete checkpoint null phenotype in taxol and nocodazole, which correlated with the absence of Mad1 at kinetochores (Fig. 1a,b and Supplementary Fig. 3a-c). We conclude that both RZZ and Bub1 have an important role in generating the checkpoint signal and that full checkpoint abrogation requires removal of both Mad1 receptors.

**Fig.1.**
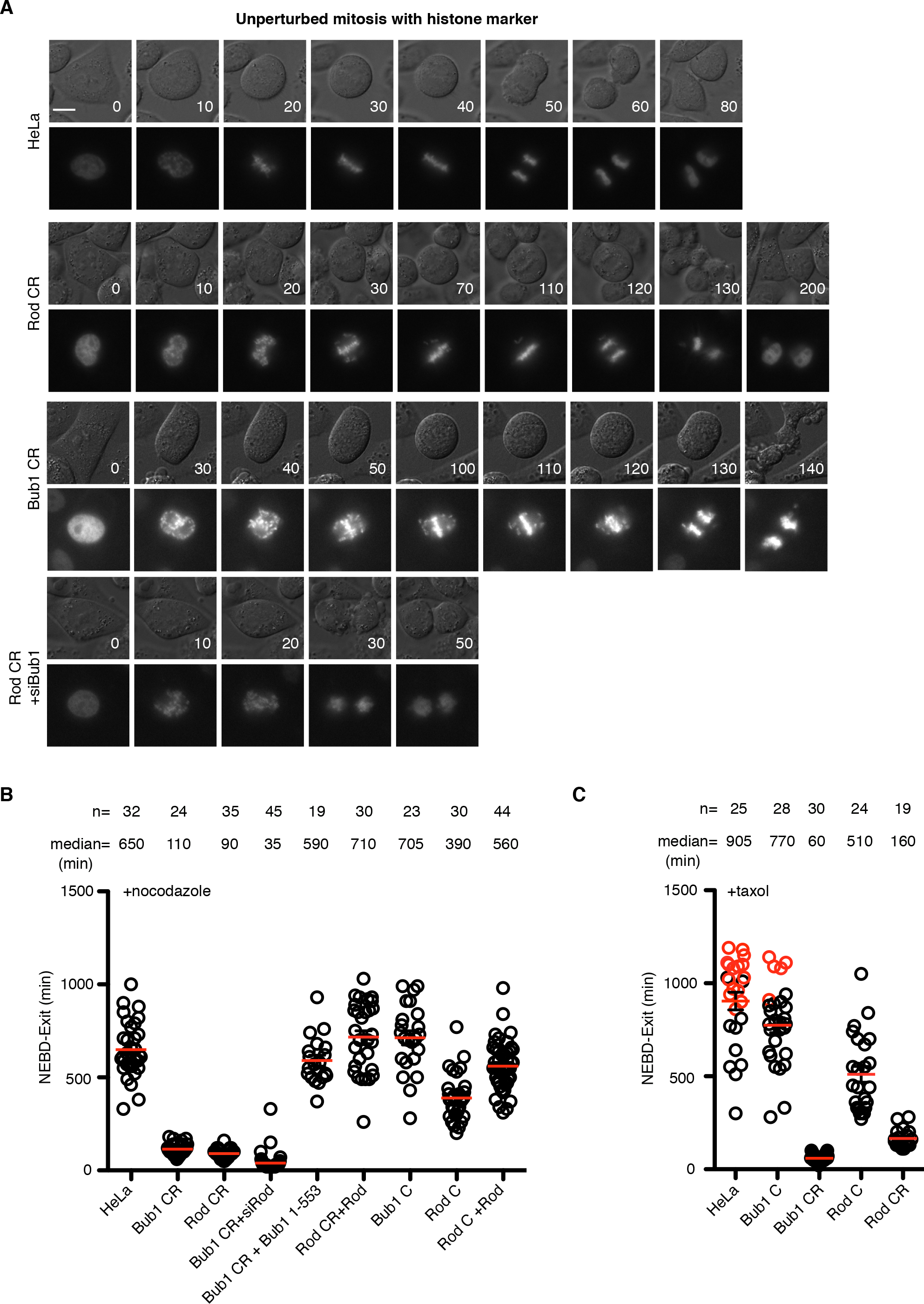
The RZZ complex and Bub1 are required for efficient checkpoint signaling. A) Representative images of indicated cell lines undergoing unperturbed mitosis with chromosomes marked by a fluorescent dye. The time from nuclear envelope breakdown (NEBD) is indicated in minutes and scale bar = 5 *μ* m. B-C) The indicated cell lines were challenged with 30 ng/ml nocodazole or 100nM taxol and analyzed by time-lapse microscopy. The time from NEBD to mitotic exit of individual cells is indicated by a circle and the median is indicated by a red line for each condition. Red circles indicates cells that were still arrested at the end of recording. The number of cells analyzed per condition (n=) and the median time are indicated above.

**Table 1.** Mitotic duration and errors in cell lines. Time from NEBD to exit in the indicated cell lines and the percent of cells delayed in alignment of chromosomes to the metaphase plate and displaying missegregation at exit.

| Condition | NEBD - exit (min) | Alignment delay(%) | Missegregation at exit (%) | n |
| --- | --- | --- | --- | --- |
| HeLa | 40 | >5 | >5 | 42 |
| Rod C | 50 | 8 | >5 | 52 |
| Roc CR | 105 | 90 | 55* | 42 |
| Bub1 C | 90 | 56 | >5 | 41 |
| Bub1 CR | 85 | 100 | 100** | 31 |
| Rod CR + Bub1 RNAi | 35 | 100 | 100 | 14 |
| Bub1 CR + Rod RNAi | 30 | 100 | 100 | 17 |

\* Few unaligned chromosomes at exit
\*\* Massive unaligned chromosomes at exit

These results raise the question as to whether Rod and Bub1 operate in separate pathways or whether they are mechanistically coupled. The quantitative comparison of Rod and Bub1 CR phenotypes suggests the latter mode of action, as their combined individual contributions to SAC strength are considerably lower than that of the control situation (Fig 1b,c). To further investigate this, we first determined if the RZZ complex is required for the interaction between Mad1 and Bub1 using our recently established proximity dependent ligation assay^9^. If Rod and Bub1 operate in separate pathways the prediction would be that depletion of Rod should not affect the Mad1-Bub1 interaction. Strikingly the removal of Rod almost completely abolished any labeling of Bub1 (Fig. 2a). This result shows that Rod is required for a full Bub1-Mad1 interaction and strongly suggests that Bub1 and Rod are functionally interlinked. Importantly the effect of Rod depletion is not an indirect effect of reduced phosphorylation of Bub1 on the S459 and T461 sites, which are required for Mad1 binding (Supplementary Fig. 1d,e).

**Fig. 2.**
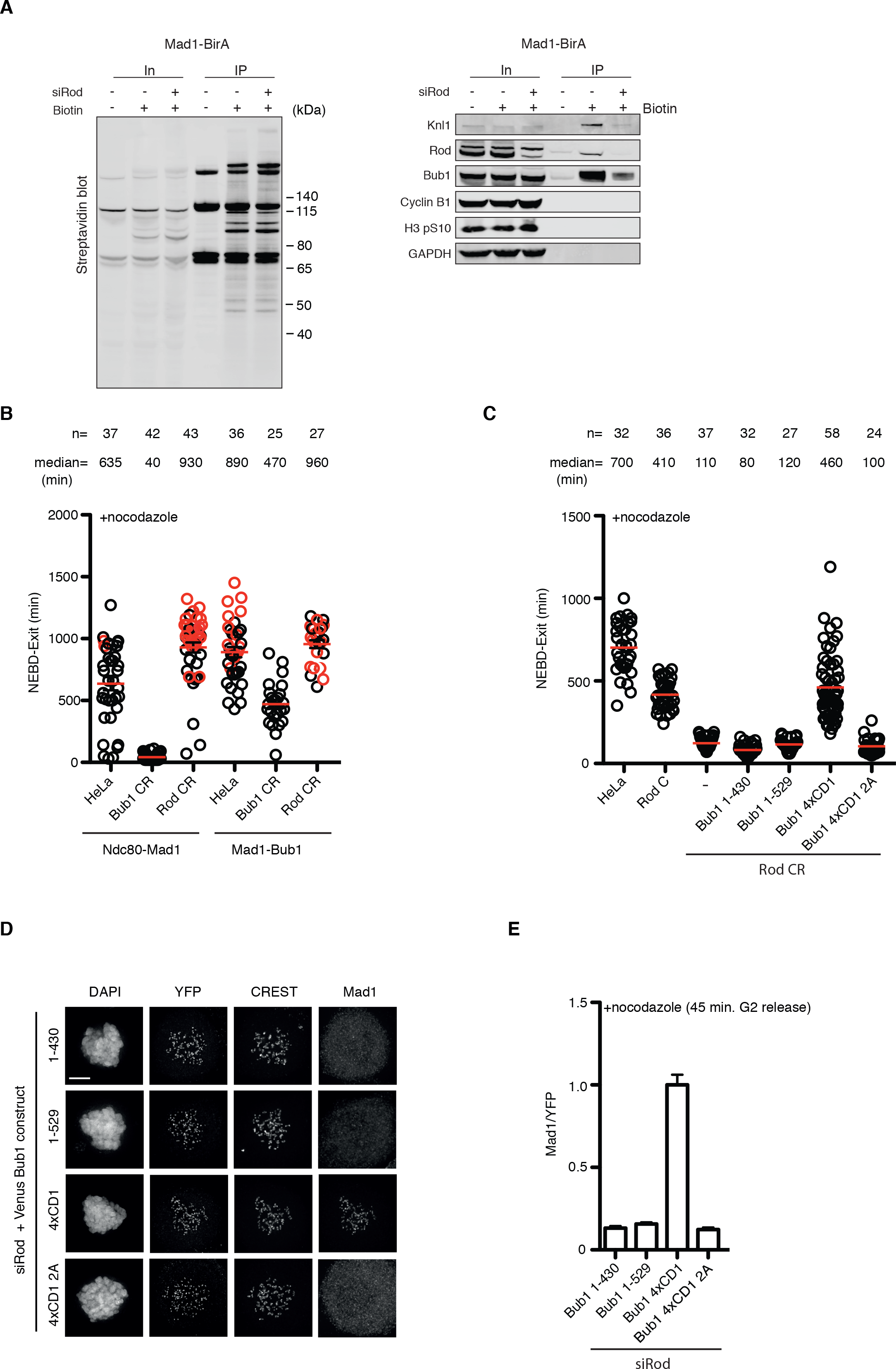
The RZZ complex stimulates Mad1 binding to Bub1. A) A HeLa cell line stably expressing Mad1-BirA were depleted of Rod and synchronized in mitosis using thymidine and nocodazole. Biotin was added to media where indicated and biotinylated proteins were purified and analyzed by western-blot with the indicated antibodies. B) Mitotic duration in the indicated cell lines expressing either Ndc80-Mad1 or Mad1-Bub1 (Mad1 485-718 fused to Bub1 1-553 lacking CD1) fusion proteins. The time from NEBD to exit is indicated and each circle represents a single cell analyzed and median indicated by red line. The red circles represents cells still arrested at the end of recording. C) Indicated cell lines expressing Bub1 fusion proteins and analyzed by time-lapse microscopy. D-E) Cells depleted of Rod using RNAi and transfected with the indicated Bub1 constructs. 45 minutes after releasing from CDK1 inhibition into nocodazole, cells were fixed and stained by YFP, CREST and Mad1 antibodies. Mad1 kinetochore levels were measured and normalized to Bub1 (YFP) level. Scale bar, 5 *μ* m.

Next, we wanted to investigate the specific role of RZZ in the checkpoint. Given the reported micromolar affinities between Mad1 and Bub1 and nanomolar concentration of these proteins in cells ^5,9,24^ we speculated that the primary function of the RZZ complex is to facilitate a high local concentration of Mad1 at kinetochores to allow productive Mad1-Bub1 interaction. If this is the case one would predict that the role of the RZZ complex in the checkpoint can be bypassed through artificial concentration of Mad1 at kinetochores. To test this we analyzed the SAC in Rod CR and Bub1 CR when Mad1 is tethered to the outer kinetochore protein Ndc80 (Fig. 2b) ^9,25^ While Bub1 is still required for efficient checkpoint signaling in Ndc80-Mad1 Rod is not. A similar result was obtained by fusion of Mad1 to the outer kinetochore protein KNL1 and using RNAi depletion (Supplementary Fig. 3d). This shows that the primary role of the RZZ complex is to recruit Mad1 to the kinetochore and further suggests that this stimulates the checkpoint generating Bub1-Mad1 interaction.

If this hypothesis is correct then enhancement of the Mad1-Bub1 interaction should be sufficient to support SAC signaling in the absence of RZZ. To test this directly we sought of ways to artificially stimulate the Mad1-Bub1 interaction. Interestingly, plants and algae lack the RZZ complex and one of their three Bub1 like proteins contains multiple repeats of the CD1 domain likely to increase the strength of the Mad1-Bub1 interaction^26,27^. To mimic this in human cells, we generated a Bub1 construct containing four repeats of the CD1 domain (Bub1-4XCD1). As a control, a similar construct but with the four CD1 domains lacking the two phosphorylation sites mediating Mad1 interaction was used (4XCD1 2A). Bub1-4XCD1 very efficiently recruited Mad1 to kinetochores and supported efficient checkpoint signaling in Rod CR cells while Bub1-4XCD1 2A or Bub1 1-529 with only one CD1 did not (Fig. 2c-e). In a complementary approach Mad1 was fused directly to Bub1 and this also bypassed the requirement for Rod (Fig. 2b and supplemental Fig.3d). Altogether these results show that Bub1 and Rod are functionally coupled and that the primary role of RZZ in the SAC is to localize Mad1 at kinetochores allowing for a functional Mad1-Bub1 interaction.

To further probe the function of Bub1 in the checkpoint, the localization of Venus-Rod to kinetochores was investigated in Bub1 depleted cells. Efficient Rod kinetochore localization was dependent on Bub1 supporting that Bub1 stimulates its own interaction with Mad1 by facilitating RZZ localization (Fig. 3a and supplemental Fig. 4a) ^28,29^. Whether there is a coupling between Mad1 binding to Bub1 and RZZ localization is unclear. To test this, we determined if the Mad1 binding and RZZ recruitment activities of Bub1 can be uncoupled. We first analyzed RZZ localization in two Bub1 mutants, Bub1 S459A/T461A (Bub1 2A) and I471D, unable to bind Mad1 (Fig. 3b-d and supplemental Fig. 4b) ^9^. In both of the Bub1 mutants RZZ localization was normal while Mad1 levels was reduced by 20-30% (Fig. 3b,c and supplemental Fig. 4c). This argues that Mad1 binding to Bub1 is unlikely to account for the function of Bub1 in stimulating recruitment of RZZ. Interestingly the Bub1 MFQ/RKK mutant, also unable to bind Mad1, strongly reduced Bub1 dependent recruitment of RZZ suggesting that a distinct region of CD1 contributes to RZZ localization (Fig. 3c,d). Furthermore the region downstream of CD1, amino acids 485-521, was required for RZZ localization which shows that these residues together with CD1 stimulates RZZ localization (Fig. 3c,d). Indeed if we fused the kinetochore targeting domain of Bub1 to a region encompassing CD1 and the downstream region this was sufficient for RZZ recruitment (Fig. 3b,e,f). In conclusion, Bub1 stimulation of RZZ localization is independent of the Bub1-Mad1 interaction but the region in Bub1 responsible for RZZ localization is partly overlapping with the Mad1 interaction domain.

**Fig. 3.**
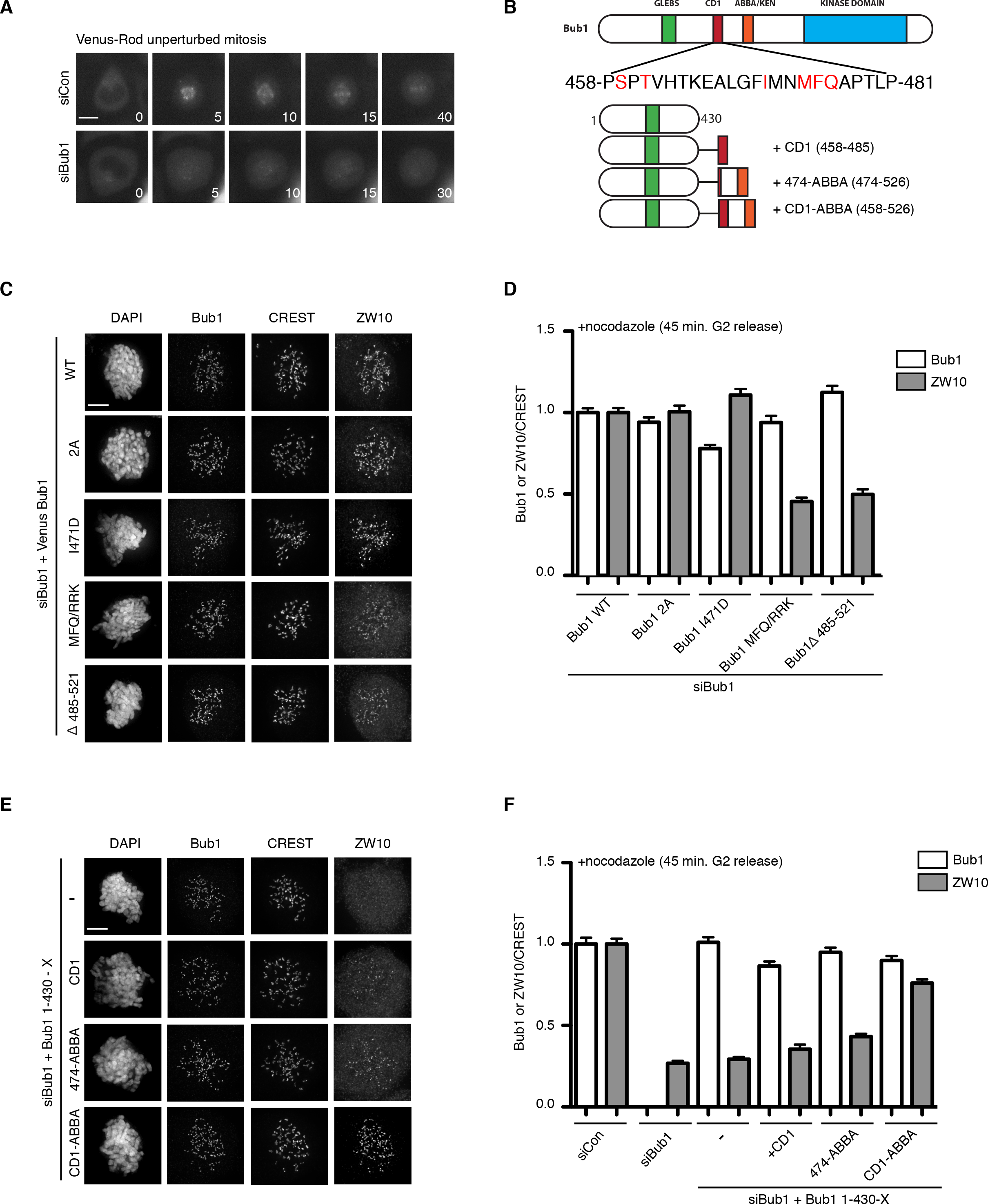
Bub1 recruitment of RZZ to kinetochores. A) Localization of Venus-Rod during unperturbed mitosis in control depleted or Bub1 depleted cells. B) Primary structure of Bub1 with sequence of CD1 shown and residues mutated in red. Schematic of fusion constructs used in panel E-F. C) HeLa cells depleted of Bub1 using RNAi and complemented with indicated Venus-Bub1 constructs. 45 minutes after releasing from CDK1
inhibition into nocodzole, cells were fixed and stained by Bub1, CREST and ZW10 antibodies. D) Quantification of Bub1 and ZW10 kinetochore levels normalized to CREST in the indicated conditions. E-F) As C-D) but with the indicated Bub1 fusion proteins.. Scale bar, 5 *μ* m.

We propose that the function of Bub1 in Mad1 recruitment is two-fold and highly integrated with the RZZ complex (Fig. 4). The central Mad1 binding region of Bub1 stimulates RZZ localization hereby ensuring efficient Mad1 kinetochore localization to facilitate its own interaction with Mad1. Since the RZZ complex localizes to kinetochores quickly after nuclear envelope breakdown we envision this mechanism is established immediately. It is important to point out that Bub1 only stimulates RZZ recruitment and if sufficient time is provided then in the absence of Bub1 the RZZ complex will localize and generate a fibrous corona that can recruit Mad1 ^17,29^. However the strong SAC defect observed in the absence of Bub1 show that RZZ bound Mad1 is less efficient in SAC signaling. Similarly Bub1 can in the absence of RZZ recruit Mad1 at reduced levels, which generates a weakened checkpoint signal. It is presently unclear to us why the Mad1 recruited by Bub1 in the absence of RZZ is not generating an efficient checkpoint signal. A possibility is that a Mad1-Bub1-RZZ complex exists and that RZZ in this complex stabilizes the Mad1-Bub1 interaction but this function of the RZZ is not needed when Mad1 is artificially tethered to kinetochores. Alternatively a gradual reduction in the Bub1 phosphorylations mediating Mad1 interaction only allows Mad1-Bub1 complex formation for a limited time in the absence of RZZ^10^. The ability of Bub1 and RZZ to independently generate a weak checkpoint signal likely explains why mammalian cells that efficiently align their chromosomes can propagate without one of the Mad1 receptors although adaption in these cell lines cannot be presently excluded ^20,30,31^. The combination of CRISPR and RNAi used here avoids issues with adaption while still obtaining penetrant depletion and robust phenotypes that can be rescued with RNAi resistant constructs. Potentially this can be a general approach to study essential genes.

**Fig. 4.**
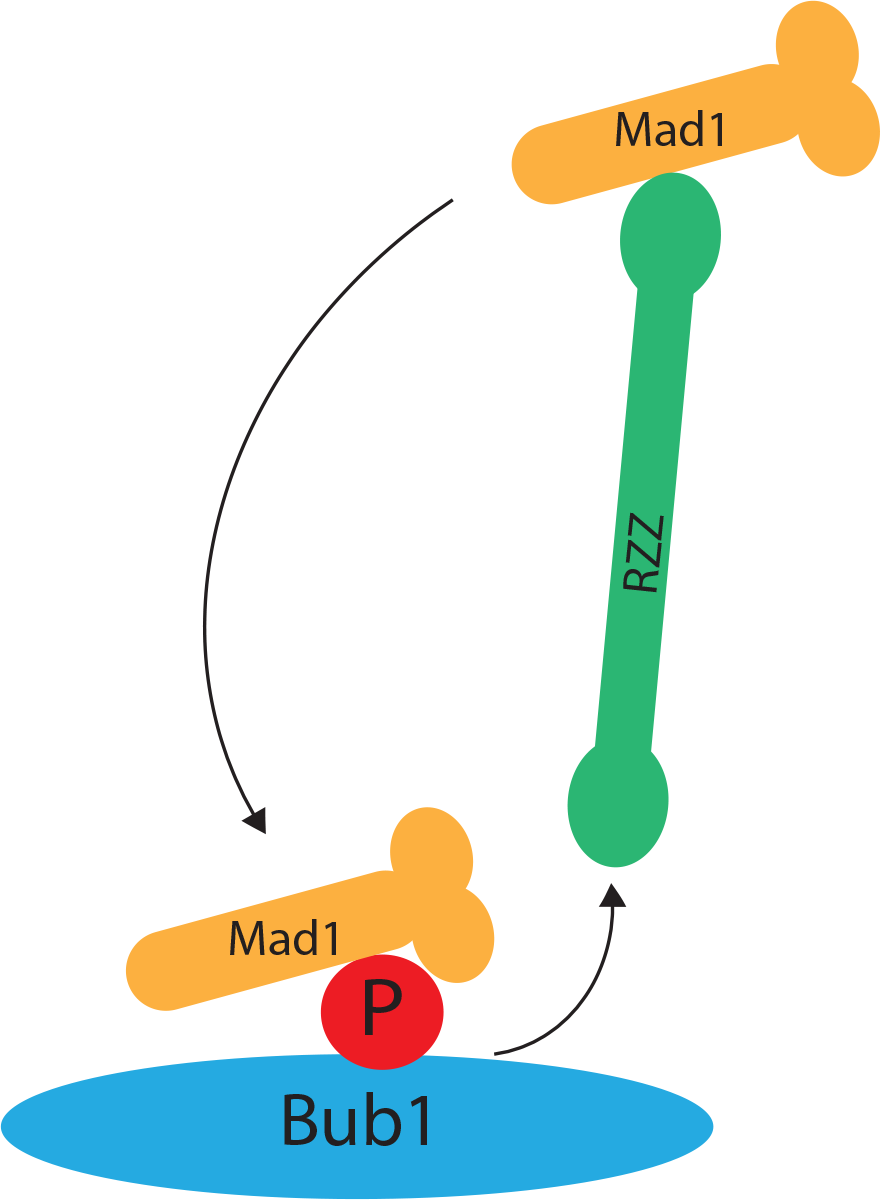
Model for the role of RZZ and Bub1 in Mad1 localization. Model depicting the role of Bub1 in stimulating RZZ recruitment and RZZ in stimulating Mad1 binding to Bub1.

We favor that checkpoint strength is largely set by the amount of Mad1-Bub1 complex at kinetochores, which is not directly reflected by Mad1 kinetochore levels. Importantly the amount of Mad1-Bub1 complex likely changes over time as the amount of MCC that needs to be produced for establishing and maintaining the SAC are different. An initial high level of Bub1 phosphorylation might ensure that a large fraction of Mad1 is bound to Bub1 at early stages of mitosis to ensure a high level of MCC production to establish the checkpoint^10^. During checkpoint maintenance less Mad1-Bub1 is needed to maintain MCC levels and the bulk of Mad1 is bound to RZZ but Bub1 is still needed for efficient SAC signaling as shown by our Mad1 tethering results.

In conclusion our work reveals that the overall conceptual architecture of the checkpoint is conserved from yeast to man and likely the unique requirement for RZZ in mammalian cells reflects the weak affinity of Mad1 for Bub1.

## Acknowledgements

We thank Gordon Chan, Reto Gassman and Stephen Taylor for providing reagents and Prasad Jallepalli, Jonathan Millar and Geert Kops for fruitful discussions.

The Novo Nordisk Foundation Center for Protein Research, University of Copenhagen, is supported financially by the Novo Nordisk Foundation (grant agreement NNF14CC0001). In addition, this work was supported by grants from the Danish Cancer Society (R72-A4351-13-S2 and R124-A7827-15-S2) a grant from the Danish Council for Independent Research (DFF-4183-00388) and a grant from the Novo Nordisk Foundation (NNF160C0022394) to J.N.

## Author contributions

G.Z. performed all experiments except the Mad1 BioID experiment performed by T.K. J.N. supervised the work and all authors contributed to data interpretation and writing the manuscript.

## Supplementary materials

Material and Methods

Supplementary figures and legends

## Supplemental figure legends

**Figure 1.**
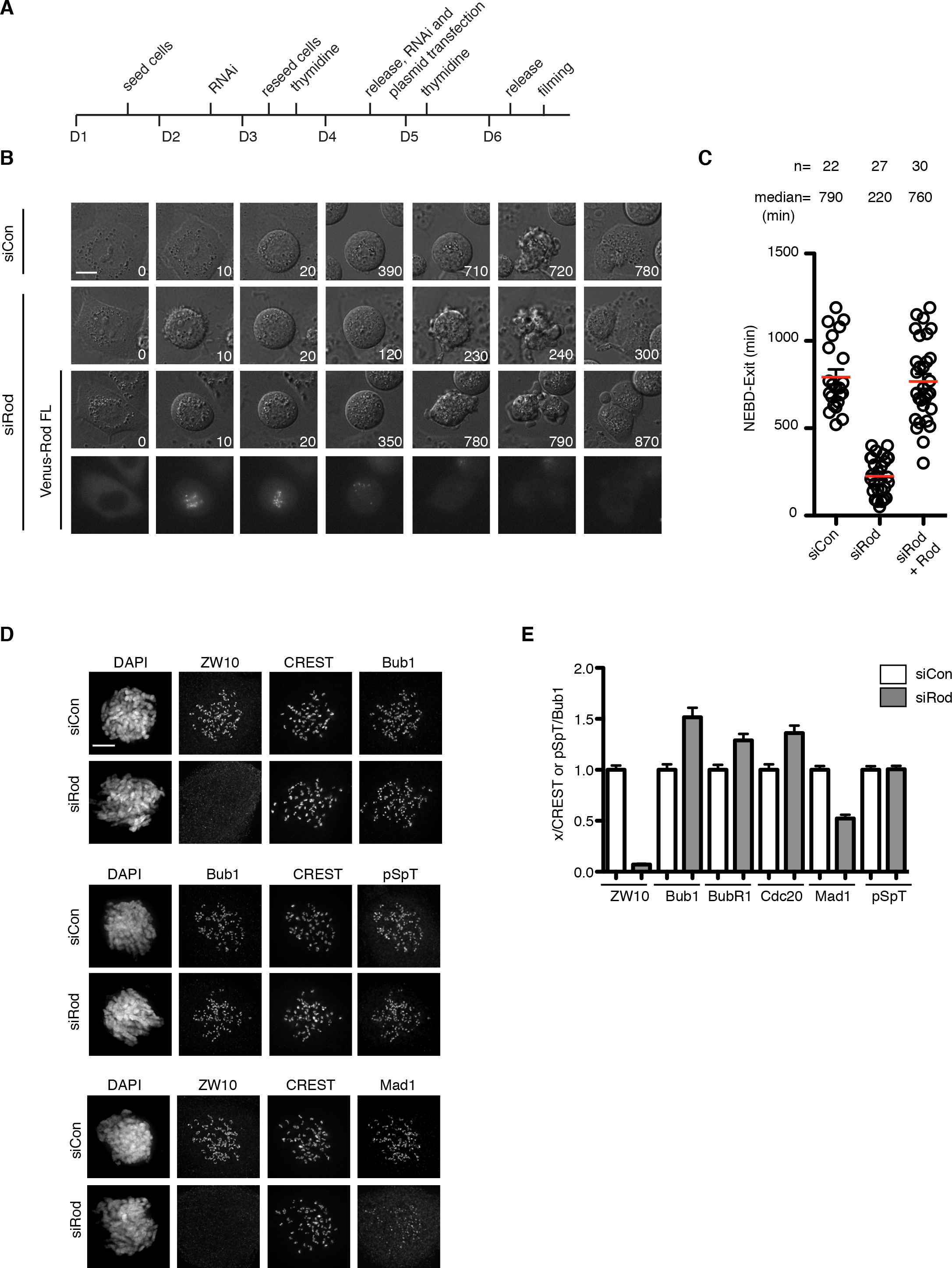
Rod is required for Mad1 localization and checkpoint signaling. A) Outline of Rod RNAi protocol and synchronization of cells. B) Representative still images from time-lapse analysis of HeLa cells arrested with nocodazole and either treated with control RNAi or Rod RNAi as indicated. An RNAi-resistant Venus-Rod construct was used to rescue the Rod RNAi phenotype as indicated. C) Quantification of NEBD-exit from time-lapse analysis of indicated conditions. Each circle represents a single cell analyzed and median time is indicated by the red line and above. Number of cells and median time are shown above. D) Representative images of immunofluoresence analysis of checkpoint proteins or Bub1 phosphorylation (pSpT) in control or Rod depleted cells. E) Quantification of kinetochore levels of the indicated checkpoint proteins normalized to CREST or Bub1 (for pSpT only). Mean and standard deviation is indicated. Scale bar, 5 *μ*m.

**Figure 2.**
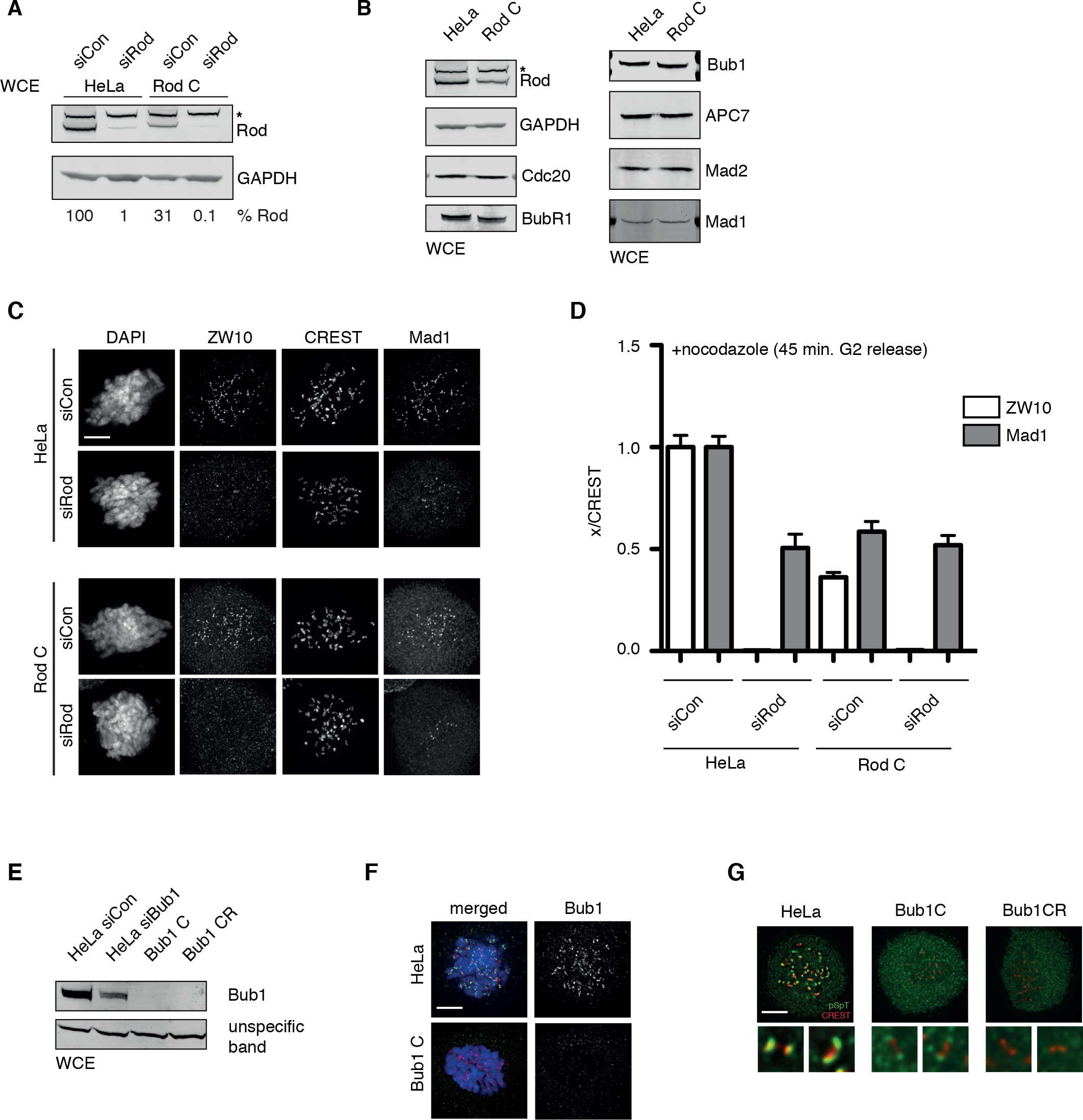
Characterization of Rod C and Bub1 C cell lines. A) Whole cell extract (WCE) from the indicated cell lines were probed for Rod and GAPDH and using quantitative LiCor technology the level of Rod was determined and indicated below in percent. * indicates unspecific band recognized by Rod antibody. B) Analysis of the protein levels of indicated checkpoint proteins in HeLa parental cell line and in Rod C cell line. C) Immunofluorescence analysis of ZW10 and Mad1 in the indicated conditions. Cells were synchronized using a thymidine block and Cdk1 inhibition and following release into nocodazole they were fixed after 45 minutes. D) Quantification of ZW10 and Mad1 levels in the indicated cell lines and conditions. The mean and standard deviation is shown. E) Western blot analysis of Bub1 in the indicated cell lines and conditions by the antibody against the N-terminus of Bub1. F) Immunofluorescence analysis of Bub1 in the indicated cell lines using an antibody against the N terminus of the protein. G) Immunofluorescence analysis of indicated cell lines with pSpT antibody recognizing Bub1 phosphorylation on Ser459 and Thr471. Green is pSpT staining while red is CREST staining. Scale bar, 5 *μ* m.

**Figure 3.**
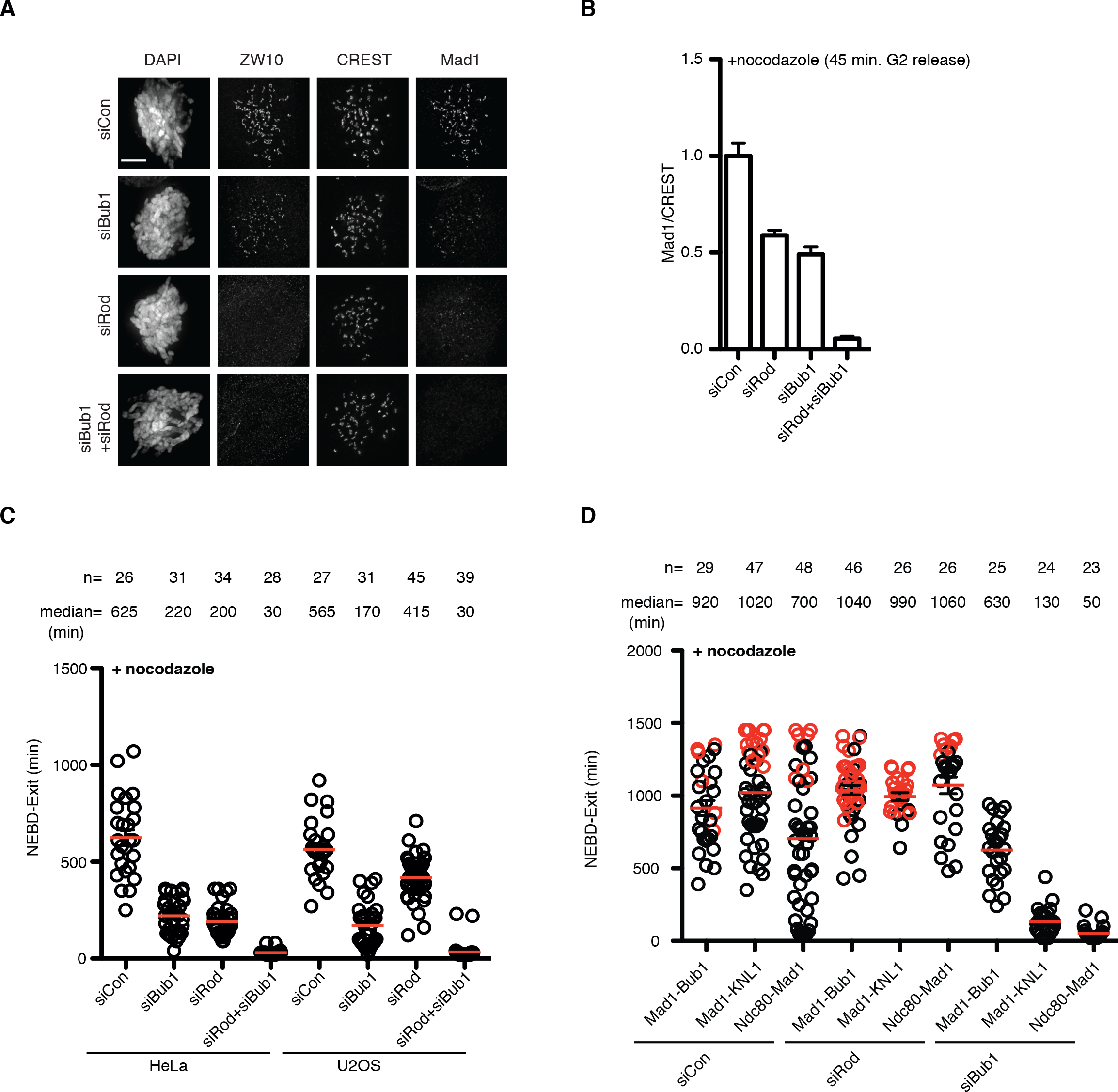
Depletion of Rod and Bub1 abrogates the checkpoint. A-B) Immunofluorescence analysis of ZW10 and Mad1 in cells treated with the indicated RNAi oligoes. Cells were synchronized with a thymidine block and Cdk1 inhibition and after 45 minutes of release from Cdk1 block into nocodazole cells were fixed and stained. The kinetochore level of Mad1 normalized to CREST is shown and control was set to 1. The mean and standard deviation is shown. C) Depletion of Bub1 and Rod in HeLa and U20S cells and quantification of mitotic duration in the presence of nocodazole. Each circle represents the time a single cell spent from NEBD to exit and a red line indicates the median. Number of cells and median time are shown above. D) Analysis of mitotic duration in HeLa cells depleted of Rod and Bub1 by RNAi and expressing the indicated Mad1 fusion constructs. Mad1-KNL1 (Mad1 485-718 fused to KNL1 1000-1200 + 1834-2316) and Mad1-Bub1 (Mad1 485-718 fused to Bub1 1-553 lacking CD1). Time from NEBD to mitotic exit is indicated by circle for single cells and median indicated by red line. Red circles represents cells still arrested at the end of recording. Number of cells and median time are shown above. Scale bar, 5 *μ* m.

**Figure 4.**
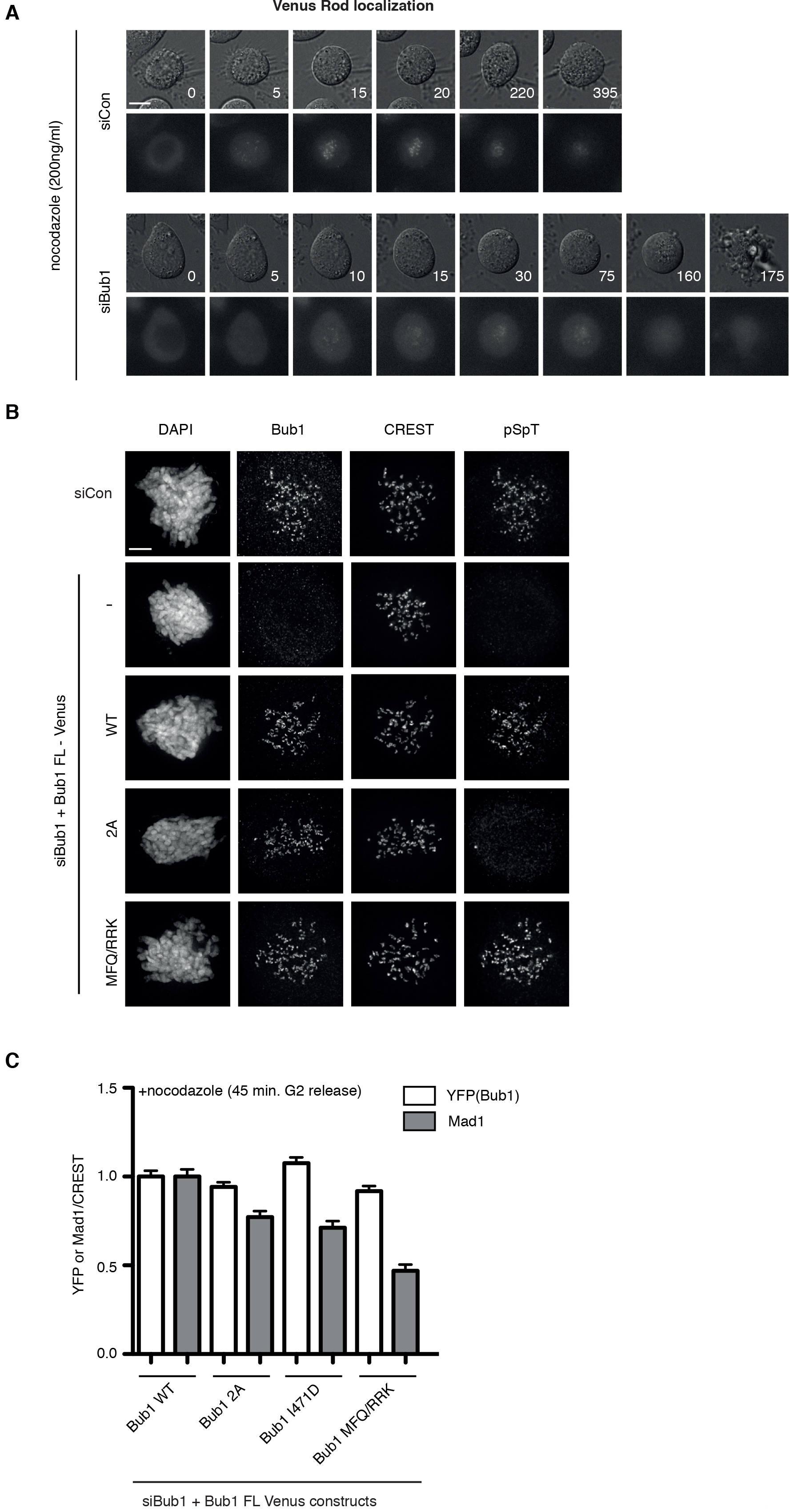
Rod localization depends on Bub1. A) Localization of Venus-Rod in control and Bub1 depleted cells treated with nocodazole. B) Immunofluorescence analysis of Bub1 phosphorylation in the indicated Bub1 constructs. Cells were depleted of Bub1 and complemented with the indicated Bub1 mutants and analyzed by immunofluorescence 45 minutes after release from a Cdk1 block into nocodazole. C) Kinetochore levels of Mad1 and YFP (Bub1 fusion protein) for the indicated Bub1 constructs normalized to CREST. The median and standard deviation is indicated. Scale bar, 5 *μ* m.

## MATERIALS AND METHODS

### Cell Culture, and RNAi

HeLa or U2OS cells were cultivated in DMEM medium (Invitrogen) supplemented with 10% fetal bovine serum and antibiotics. For Bub1 or Rod RNAi and rescue experiments, two times of RNAi were performed using RNAi Max (Invitrogen) according to manufactures instructions. The first Bub1 RNAi was performed 48 hours before filming or fixation with the second at 24 hours before in the presence of thymidine (2.5mM). The first Rod RNAi was performed 96 hours before filming or fixation with the second at 48 hours before. Thymidine was used afterwards to synchronize cells. Plasmid co-transfection was done in the first transfection for Bub1 and in the second transfection for Rod. For Bub1 and Rod co-depletion, the first Bub1 RNAi was performed at the same time as the second Rod RNAi at 48 hours before filming or fixation. RNAi oligos targeting Bub1 (5′ GAGUGAUCACGAUUUCUAA 3′) or luciferase (5′ CGUACGCGGAAUACUUCGA 3′) (Sigma) or Rod (5’ GGAAUGAUAUUGAGCUGCUAACAAA 3’) (Thermofisher), were used for RNAi depletions.

### Immunofluorescence and quantification

Cells growing on coverslips were synchronized with a double thymidine block followed by RO3306 (10 nM) block. After washing with PBS, the cells were treated with nocodazole (200 ng/ml) for 45 minutes and fixed with 4% paraformaldehyde in PHEM buffer (60 mM PIPES, 25 mM HEPES, pH 6.9, 10 mM EGTA, 4 mM MgSO_4_) for 20 minutes at room temperature. Fixed cells were extracted with 0.5% Triton X-100 in PHEM buffer for 10 minutes. The antibodies used for cell staining include Bub1 (Abcam, ab54893, 1:400), Bub1 pSpT (home made, 1:200), Rod (gift from Gordon Chan), CREST (Antibodies Incorporated, 15-234, 1:400), BubR1 (homemade), Cdc20 (Millipore, MAB3775, 1:200), ZW10 (Abcam, ab21582, 1:200), Mad1 (Santa Cruz, sc65494, 1:200). All the fluorescent secondary antibodies are Alexa Fluor Dyes (Invitrogen, 1:1000). Z-stacks 200 nm apart were recorded on a Deltavision Elite microscope (GE Healthcare) using a 100X oil objective followed by deconvolution using Softworx prior to quantification. Protein intensity on kinetochores was quantified by drawing a circle closely along the rod-like CREST staining covering the interested outer kinetochore protein staining on both ends. The intensity values from the peak three continuous stacks were subtracted of the background from neighboring areas and averaged. The combined intensity was normalized against the combined CREST fluorescent intensity. At least 100 pairs of kinetochores from 10 cells were measured per condition.

### CRISPR/Cas mediated gene knock out

GeneArt Plantinum Cas9 Nuclease (Thermofisher) with synthesized gRNA was used for gene editing. Pre-designed DNA primers for gRNA template assembly were purchased (Thermofisher) with forward primer IVT-TAATACGACTCACTATAGTACAAGGGCAATGACC and reverse primer IVT-TTCTAGCTCTAAAACAGAGGGTCATTGCCCTTGT for Bub1; with forward primer IVT-TAATACGACTCACTATAGCTTTTCTTGAACCGAC and reverse primer IVT-TTCTAGCTCTAAAACGAGTGTCGGTTCAAGAAAA for Rod. Guide RNA was synthesized according to the instruction from GeneArt Precisiion gRNA Synthesis Kit (Thermofisher). 625 ng of gRNA and 2500 ng of Cas9 nuclease (GeneArt Platinum Cas9 Nuclease, Thermofisher) were transfected into 4.5 × 10^5^ HeLa cells by Lipofectamine CRISPRMAX transfection reagent. 24 hours later, the cells were diluted and re-cultivated for single colony isolation. 20-100 single colonies were isolated and amplified followed by examination of the kinetochore signals of interested protein by immunofluorescence. The ones with reduced signals compared to the parental HeLa cells were maintained for further characterization.

### Cloning

Rod cDNA was purchased from Kazusa DNA Res Inst (ORK05777). The internal BamHI site was eliminated by Gibson Assembly and the cDNA was cloned into pcDNA5/FRT/TO N-Venus vector by BamHI and NotI sites. Venus-Bub1 1-430 and its variants were cloned into the same vector by KpnI and NotI sites. 4xCD1 WT or 2A DNA was synthesized by GeneArt (Thermofisher). Details will be provided upon request.

### Live cell imaging

Live cell imaging was performed on a Deltavision Elite system using a 40 x oil objective (GE Healthcare). Cells were transfected in 6-well plate and re-seeded in 8- well Ibidi dishes (Ibidi) one day before the filming. Growth media was changed to Leibovitz’s L-15 (Life technologies) before filming started. Appropriate channels were recorded for 18-24 hours and data was analyzed using Softworx (GE Healthcare). Statistical analysis was done using Prism software.

### Western blot analysis of cell lines

Mitotic cells induced by nocodazole (200ng/ml) were collected and lysed in lysis buffer containing 10 mM Tris pH7.5, 150 mM NaCl, 0.5 mM EDTA and 0.5% NP40. After centrifugation at 13,000 rpm for 10 min, the supernant was applied to SDS-PAGE followed by western blot with interested antibodies. The antibodies used in this study include APC7 (Bethyl, A302-551A, 1:2000), Bub1 (abcam, ab54893, 1:1000), BubR1 (home made), Cdc20 (Santa Cruz, sc13162, 1:1000), GAPDH (Santa Cruz, sc25778, 1:2000), Mad1 (Sigma, M8069, 1:1000), Mad2 (Bethyl, A300-301A, 1:1000), Rod (gift from Reto Gassmann). Fluorescent labelled secondary antibodies (1:5000, LI-COR Biosciences) were used for quantitative analysis by LI-COR Odyssey imaging system (LI-COR Biosciences).

### Purification of biotinylated protein complexes

Stable HeLa cell lines expressing the Mad1 BirA fusion protein were exposed to 0.1 ng ml^−1^ doxycycline for 18 h to obtain near endogenous Mad1 expression levels. Cells were arrested in mitosis by a double thymidine block and subsequent nocodazole (150 ng ml^−1^) treatment for 12 h. Biotinylation of proximity interactors was induced by the addition of a final concentration of 25 μM of biotin simultaneously with the addition of nocodazole. Rod siRNA knock down was performed as described above. Mitotic cells were collected and washed three times in PBS before lysed in RIPA buffer (50 mM Tris pH 7.5, 150 mM NaCl, 1 mM EDTA, 1% Nonidet P-40, 0.25% Na-deoxycholate, 0.1% SDS) containing protease inhibitors (Roche). Cell lysate was clarified by centrifugation and incubated overnight at 4 °C with High Capacity Streptavidin Resin (Thermo Scientific). Streptavidin beads were washed once with RIPA buffer followed by two washes with water containing 2% SDS and a final wash with RIPA buffer. Biotinylated proteins were eluted from the streptavidin beads with 2 × Laemmli LDS sample buffer containing 1 mM of biotin before separated on 4-12% Bis-Tris NuPage gels (Life technologies). After separation, proteins were examined by western blot using following antibodies: Cyclin B1 (554177, 1:1000, BD Pharmingen, H3 pS10 (06-570, 1:1000, Millipore), GAPDH (sc-25778, 1:500, Santa Cruz Biotech.), Bub1 (ab54893, 1:1000, abcam) Knl1 (produced in house, 1:1000), Rod (1:2000, gift from Reto Gassmann).

